# *Saccharomyces cerevisiae* strains display robust phenotypes in the presence of Dyskeratosis congenita mutations in the *Cbf5* gene

**DOI:** 10.1101/535443

**Authors:** Abeer Abdullah Ogailan, Anne C. Rintala-Dempsey, Ute Kothe

## Abstract

Dyskeratosis congenita is a rare, congenital disorder affecting the skin, nails and oral mucosa of patients that often progresses to bone marrow failure and an increased predisposition for a variety of carcinomas. Mutations in the human dyskerin gene have been identified as the most prevalent cause of the disease. Dyskerin is a pseudouridine synthase and the catalytic subunit of H/ACA ribonucleoproteins (RNPs) responsible for the modification of uridines to pseudouridine in ribosomal RNA (rRNA), but dyskerin also binds to the telomerase RNA component (TERC). Accordingly, Dyskeratosis congenita mutations have been reported to affect both telomerase function as well as ribosome biogenesis, but the relative contribution of each pathway to the diseases is under debate. As the yeast homolog of dyskerin, Cbf5, does not interact with telomerase RNA, *Saccharomyces cerevisiae* is an ideal model to identify the selective impact of Dyskeratosis congenita mutations on ribosome biogenesis. Therefore, chromosomal mutations in the yeast homologue of dyskerin, Cbf5, were introduced at positions corresponding to the mutations in human dyskerin that result in Dyskeratosis congenita. To determine if the mutations affect cellular fitness, we screened for growth defects in yeast. Growth curves at different temperatures and yeast spot assays under several stress conditions revealed that the mutations in *cbf5* did not impair growth compared to wild type. These findings suggest that in the yeast cell, Dyskeratosis congenita mutations do not significantly affect ribosome biogenesis, and we discuss the implications for understanding the molecular cause of Dyskeratosis congenita.

## Introduction

Dyskeratosis congenita, also known as Zinsser-Cole-Engman syndrome, is a rare congenital disorder that was first observed by Zinsser in 1910 [1]. Symptoms and severity vary widely among affected individuals, but usually include the triad of abnormal skin pigmentation, nail dystrophy and leukoplakia of oral mucous membranes [2]. The disease often leads to aplastic anemia, or bone marrow failure, due to reduced red blood cell production which is the primary cause of premature death. In addition, patients also have an increased risk of developing leukemia and other cancers as well as pulmonary fibrosis resulting in reduced oxygen transport [3]. Originally thought to be X-linked and affecting only males, Dyskeratosis congenita was later found to also affect a smaller percentage of females and could either be autosomal recessive or dominant depending on the affected gene [4, 5].

Dyskeratosis congenita is caused by mutations in several different genes (namely *DCK1, TERT, TERC, TINF2, NOP10, NHP2, ACD, WRD79, CTC1* and *RTEL1*), all of which are involved in chromosome maintenance [6, 7]. In brief, *TERT* codes for the protein component of the telomerase complex and is responsible for adding telomere segments to the ends of chromosomes, while *TERC* codes for the RNA component of the complex, called hTR. The *TINF2, RTEL1*, and *ACD* genes code for proteins that are members of the shelterin complex and involved in the protection of the telomere. The CTC1 protein is a member of CST complex involved in telomere maintenance during stress conditions while *WRD79* codes for TCAB1, a protein responsible for the localization of the telomerase complex to Cajal bodies.

Mutations in the *DKC1* gene result in the most abundant and also most severe cases of Dyskeratosis congenita [8–10]. *DKC1* codes for dyskerin, a protein with multiple functions. On the one hand, dyskerin binds to the hTR RNA stabilizing the telomerase complex [11, 12]. On the other hand, dyskerin is also a critical subunit of H/ACA small nucleolar ribonucleoproteins (snoRNPs) [13]. This family of enzymes is responsible for the modification of specific uridines to generate all pseudouridines in rRNA, but also introduces some pseudouridines in small nuclear RNA (snRNA), messenger RNA (mRNA) and other RNAs [14, 15]. Dyskerin is the catalytic subunit of the ribonucleoprotein complex whereas the guide RNA directs the H/ACA snoRNP to the target uridine by base-pairing interactions [16]. Moreover, at least one specialized and essential H/ACA RNA in complex with dyskerin is required for processing of rRNA [17]. Over 50 mutations in the *DKC1* gene have been identified as causes of Dyskeratosis congenita [10]. The majority of the mutations result in single amino acid substitutions across all domains of the dyskerin protein including the catalytic domain, the PUA domain, a positive amino acid rich region binding RNA, and the N- and C-termini.

Given the different functions of dyskerin, it remains unclear to which extent impaired ribosome biogenesis contributes to the development of Dyskeratosis congenita which is often considered to be a telomere-linked disorder [10, 18, 19]. In this context, it is notable that the yeast homologue of human dyskerin called Cbf5 is the only essential pseudouridine synthase in yeast and is the sole enzyme responsible for all pseudouridines found in rRNA. Together with the H/ACA guide RNA snR30, Cbf5 also contributes to rRNA processing. However, unlike in mammals, the telomerase RNA in yeast does not interact with Cbf5 or the other H/ACA proteins [20]. Therefore, this study aimed at dissecting the impact of Dyskeratosis congenita mutations in *cbf5* on ribosome biogenesis without confounding effects on telomere maintenance by generating *S. cerevisiae* model strains.

## Materials and methods

### Materials

All chemicals and biochemical were purchased from Fisher Scientific unless noted otherwise. Yeast media was obtained from Sunrise. Oligonucleotides were from Integrated DNA Technologies (IDT).

### Preparation of Yeast-integrating plasmids harboring *cbf5* mutations

Overlapping primers for each *cbf5* mutation were used to introduce mutations in a pUC19-ScCbf5 plasmid by QuikChange mutagenesis using Q5 DNA polymerase (NEB) (Table 1). The pUC19-ScCbf5 plasmid contains the entire *CBF5* coding region flanked by 33 nt upstream and 348 nt downstream of the coding region. After DpnI digestion, the mutagenesis products were transformed into *E. coli* DH5α cells. Subsequently, the mutated *cbf5* gene was excised from pUC19-ScCbf5 by EcoRI and SalI restriction and ligated into equally digested and dephosphorylated YIp5 plasmid and transformed into *E. coli* DH5α competent cells. All plasmid sequences were confirmed by sequencing (Genewiz).

**Table 1:**
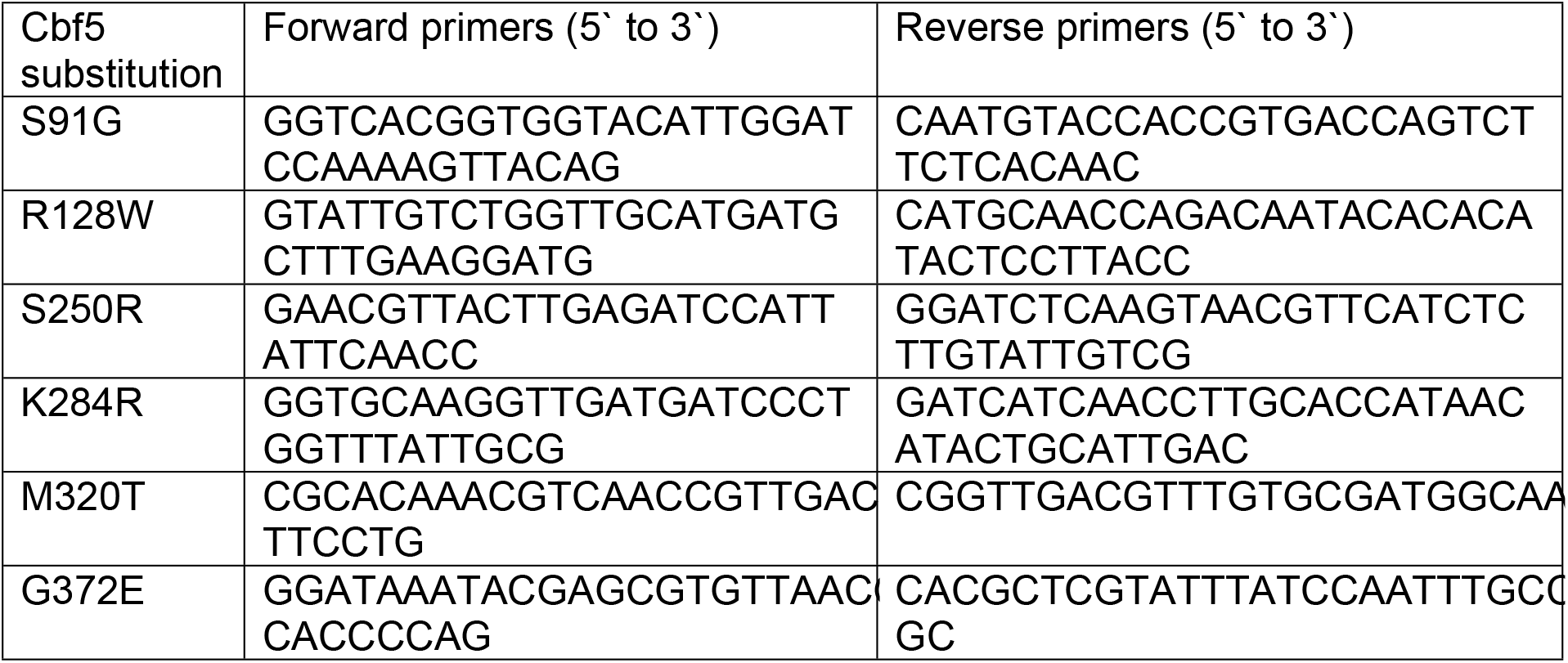
Overlapping primers to introduce mutations into *Cbf5*. The resulting amino acid substitution in the *S. cerevisiae* Cbf5 protein is stated.

### Generation of mutant yeast strains

The YIp5-*Sccbf5(mutant*) plasmids were linearized by single restriction within the *cbf5* gene using BshTI for *cbf5* K284R, M320T and G372E or HpaI (KspAI) for *cbf5* S91G and R128W and BamHI for S250R. The linearized YIp5-*Sccbf5* plasmids were transformed into haploid BY4741 cells and plated onto Sc-ura + 2% glucose plates resulting in integration of the plasmid within the chromosomal *CBF5* gene [21, 22]. The transformant had one full-length *cbf5* gene followed by the YIp5 sequence and an additional copy of the coding region lacking a promoter. After transformation, genomic DNA from yeasts cells was extracted by using Geneaid-Presto Mini gDNA Yeast Kit from FroggaBio Scientific Solutions following the manufacturer’s instructions. To verify the correct integration of the YIp5-*Sccbf5* plasmid and the presence of the mutation, a PCR reaction of genomic DNA was prepared using the forward primer EcoR1-ScCbf5-up and the reverse primer YIp5 downstream SalI reverse (Table 2). The PCR product was visualized via agarose gel electrophoresis and confirmed by sequencing (Genewiz, see Table 2 for sequencing primers). The location of the primers relative to the chromosomal *cbf5* gene is illustrated in Fig 1A.

**Table 2:**
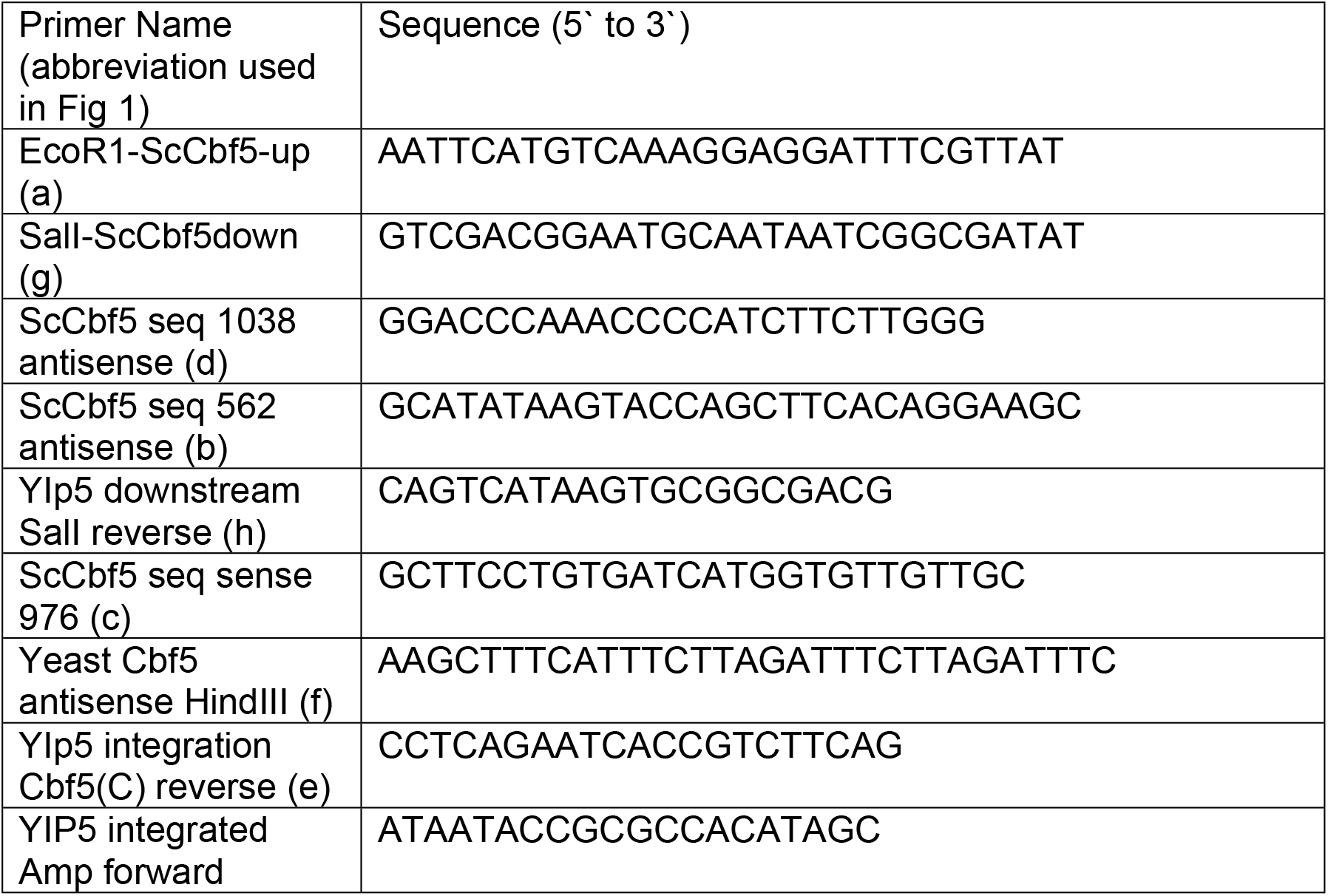
Primers used to amplify *cbf5* or for sequencing.

**Fig 1:**
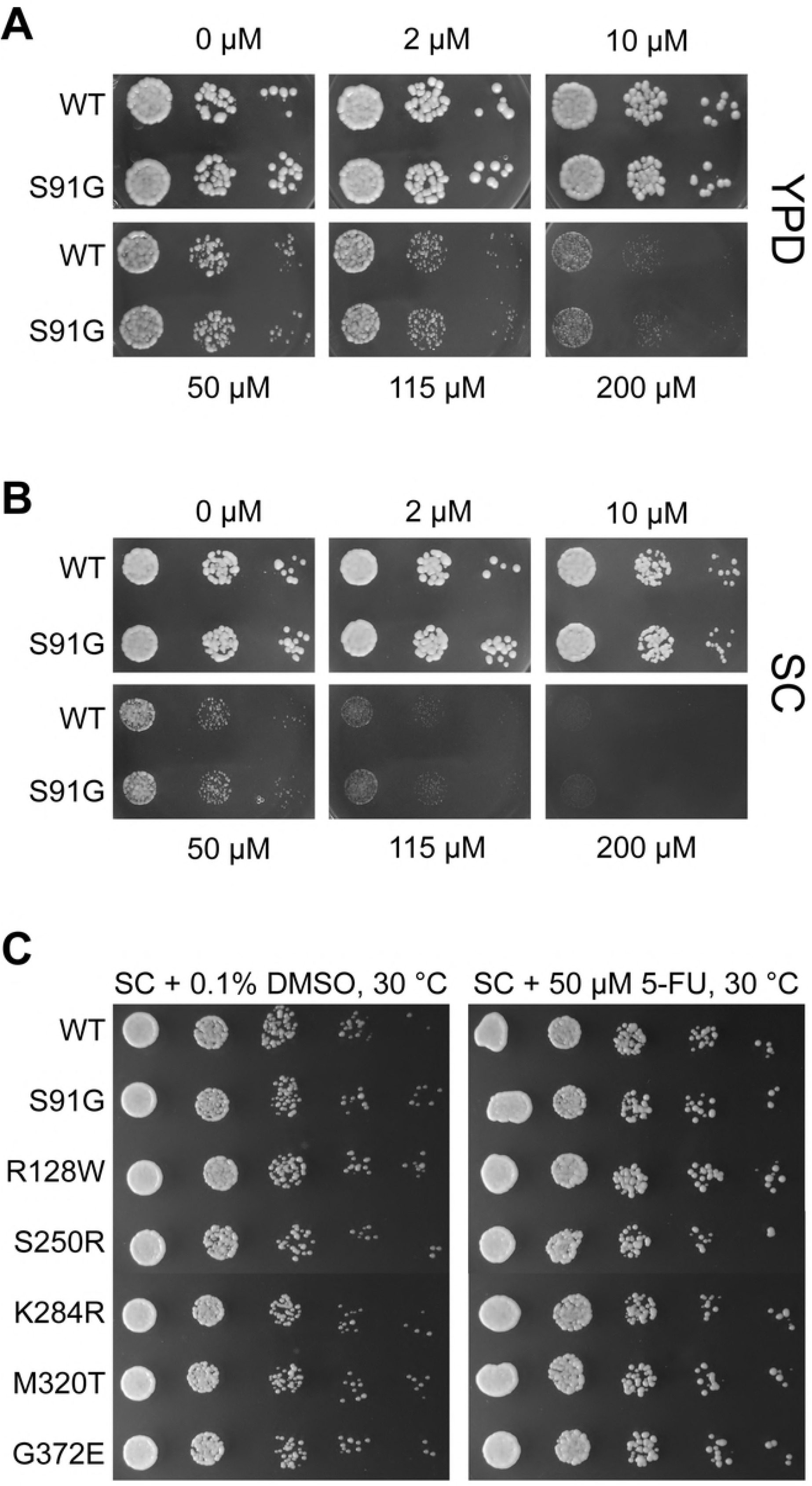
Location of Dyskeratosis congenita substitutions in *S. cerevisiae cbf5* used in this study. (A) Linear representation of the Cbf5 protein sequence including domain representation. The catalytic domain of Cbf5 is shown in blue and the PUA domain in magenta with amino acid substitutions indicated above. Primers used in the generation of the mutant strains are shown below (see Table 2). (B) Mapping of amino acid substitutions onto the crystal structure of yeast Cbf5-Nop10-Gar1 (PDBID: 3U28). The substitutions S91G, R128W and S250R are located in the catalytic domain whereas the substitutions K284R and M320T are in the PUA domain. The G372E mutation is within the unresolved C-terminal region and is not shown.

To remove the YIp5 plasmid sequence and the additional *cbf5* coding region from the genome, yeast strains were streaked on SD-ura plates, and one colony was selected and incubated in 5 mL YEPD liquid culture at 30°C for 7 to 8 h until the cells reached the exponential phase (0.4 OD_600_). The cells were collected via centrifugation at 3000 *xg* for 5 min and resuspended in YEPD to 5×10^8^ cells/mL. Serial dilutions of 1×10^6^ and 1×10^7^ cells/mL were plated on 5.7 mM 5-fluoroorotic acid (5-FOA) plates. The plates were incubated at 30°C for 48 h. Following the plasmid removal, genomic DNA was prepared of selected colonies as described above, the *cbf5* gene was amplified using the primers “EcoR1-ScCbf5-up” and “YIp5 downstream SalI reverse”, and the PCR products were sequenced to identify strains harboring the *cbf5* mutation.

### Growth analysis in liquid medium

All yeast cells were grown in liquid YPD + 2% glucose unless otherwise specified. Three colonies (biological replicates) from each strain were selected from YPD plates and grown at 30°C and 200 rpm shaking for 5 hours. For each culture, three technical replicates were generated by preparing three dilutions of 1×10^6^ of cells/130 μL in growth medium. Each dilution was placed into one well of a 96-well plate; as blank, YPDE+ 2% glucose was used. The OD_600_ was measured every 15 min in a BioTeK microplate reader with Gene5 software. This experiment was conducted at 30°C, 18°C and at 37°C with shaking at 250 rpm. The average growth curve of the three technical replicates was calculated.

### Yeast Spot Assays

Single colonies were used to inoculate culture tubes containing 5 mL YPD with 2% glucose and grown for approximately 16 h at 30 °C with shaking. Dilutions of each culture in sterile water were prepared to 0.5 OD_600_ into a 96-well plate. Serial 1:10 dilutions were performed in the 96-well plate using sterile water. Spots of 5 μL each were pipetted onto either YPD or SC agar plates. Various stress conditions were tested by preparing YPD and SC plates containing glycerol, ethanol or formamide. YPD agar with 2% glucose (except when glycerol was added) was supplemented with final concentrations (v/v) of 3% glycerol, 4% ethanol or 3% formamide. Spot tests were performed in duplicate.

For the 5-fluorouracil yeast spot test, a 50 mM stock solution of 5-fluorouracil was prepared in DMSO. To determine the concentration of 5-fluorouracil to use in yeast spot assays, 6-well plates with agar containing 0, 5, 10, 50, 115 or 200 μM 5-fluorouracil (BioBasic) in each well were used [23]. The 0 μM agar contained DMSO as a control. Each well was spotted with 4 μL each of three serial dilutions of both wild-type BY7471 cells and the cbf5-S91G cells. Plates were incubated for 2 days at 30 °C. For the testing of all strains, a final concentration of 50 μM 5-fluorouracil (1:1000 dilution) was used. In addition, control SC plates with 0.1% DMSO were also prepared.

### Western blotting

Whole-cell extracts were prepared from 2 OD_600_ of cells from exponentially growing cultures. After harvesting, the cells were resuspended in 0.4 mM NaOH for 5 min. After a second centrifugation, the cells were resuspended in 0.1 M Tris pH8, 8 M Urea, 10% SDS and boiled for 7 min before adding 6x SDS loading dye and additional boiling for 3 min. The total protein concentration was determined by a Bradford assay. To detect and semi-quantify Cbf5 protein levels, whole-cell extracts were separated by 12% SDS-PAGE and blotted onto nictrocellulose. First, the membrane was incubated with 1:1000 PGK1 primary antibody conjugated with horseradish peroxidase (Abcam) overnight as a loading control. The membrane underwent a mild stripping procedure and was then probed for Cbf5. Primary antibody against *S. cerevisiae* Cbf5 was custom-made from BioBasic Inc. by infecting two rabbits with an antigen of a 15-residue long, synthesized peptide designed from a sequence at the C terminus of Cbf5 (aa 461-475: KKEKKRKSEDGDSEE). The blot was incubated with a 1:1,000 dilution in 3% BSA in TBS of this primary antibody overnight at 4°C. The next day, the blot was exposed to a secondary antibody (horseradish peroxidase coupled anti-rabbit antibody (Sigma), 1:1,000 dilution in 3% BSA and 1x TBS), and the bands were visualized by chemiluminescence.

## Results

To determine how mutations in *cbf5* affect the ribosome, several human disease-causing mutations in dyskerin were introduced at the corresponding sites in the yeast *cbf5* gene (Fig 1). Representative mutations were chosen from different regions of the protein [24]. Specifically, point mutations in the dyskerin gene that result in the disease-causing amino acid substitutions S121G, R158W, S280R, K314R, M350T and G402E were chosen for mutation in yeast *cbf5*. The corresponding mutations in the Cbf5 protein are S91G, R128W and S250R in the catalytic domain, K284R and M320T in the PUA domain and G372E in the C-terminal region. Notably, the S121G substitution is found in patients suffering from the severe Dyskeratosis congenita variant called Hoyeraal-Hreidarsson syndrome, and the K314R and the M350T substitutions are present in multiple families [10]. Each single mutation was introduced into the yeast genome and confirmed by sequencing [22].

### Growth Assays

To determine if the mutations in *cbf5* result in growth defects in yeast cells by possibly affecting ribosome biogenesis, we first conducted growth assays in liquid medium at different temperatures. Sensitivity of yeast cells to heat often indicates the mutation is reducing protein stability while cold sensitivity can suggest problems with subunit assembly [25]. Growth curves were recorded at 18, 30 and 37 °C in a plate reader until cells reached stationary phase (Fig 2). At 30 °C, none of the mutant yeast strains is growing significantly slower than the wild-type strain (Figs 2 A and B); notably some mutations seem to enable faster growth, in particular for *cbf5-S250R, cbf5-M320T* and *cbf5-G372E* cells. Next, the cells were grown at low temperature of 18 °C to determine if any of the *cbf5* mutants displayed cold sensitivity (Figs 2 C and D). Again, all mutant yeast strains display a growth curve that is overall similar to wild-type. For a high temperature growth assay at 37 °C (Figs 2 E and F), similar results were obtained as no mutant yeast strain has impaired growth relative to the wild-type. But again, the *cbf5-S250R* and the *cbf5-M320T* cells appear to grow slightly faster than wild-type as also observed at 30 °C. These experiments show that in liquid medium, the mutations in *cbf5* do not significantly impair yeast growth and do not cause heat- or cold-sensitivity.

**Fig 2:**
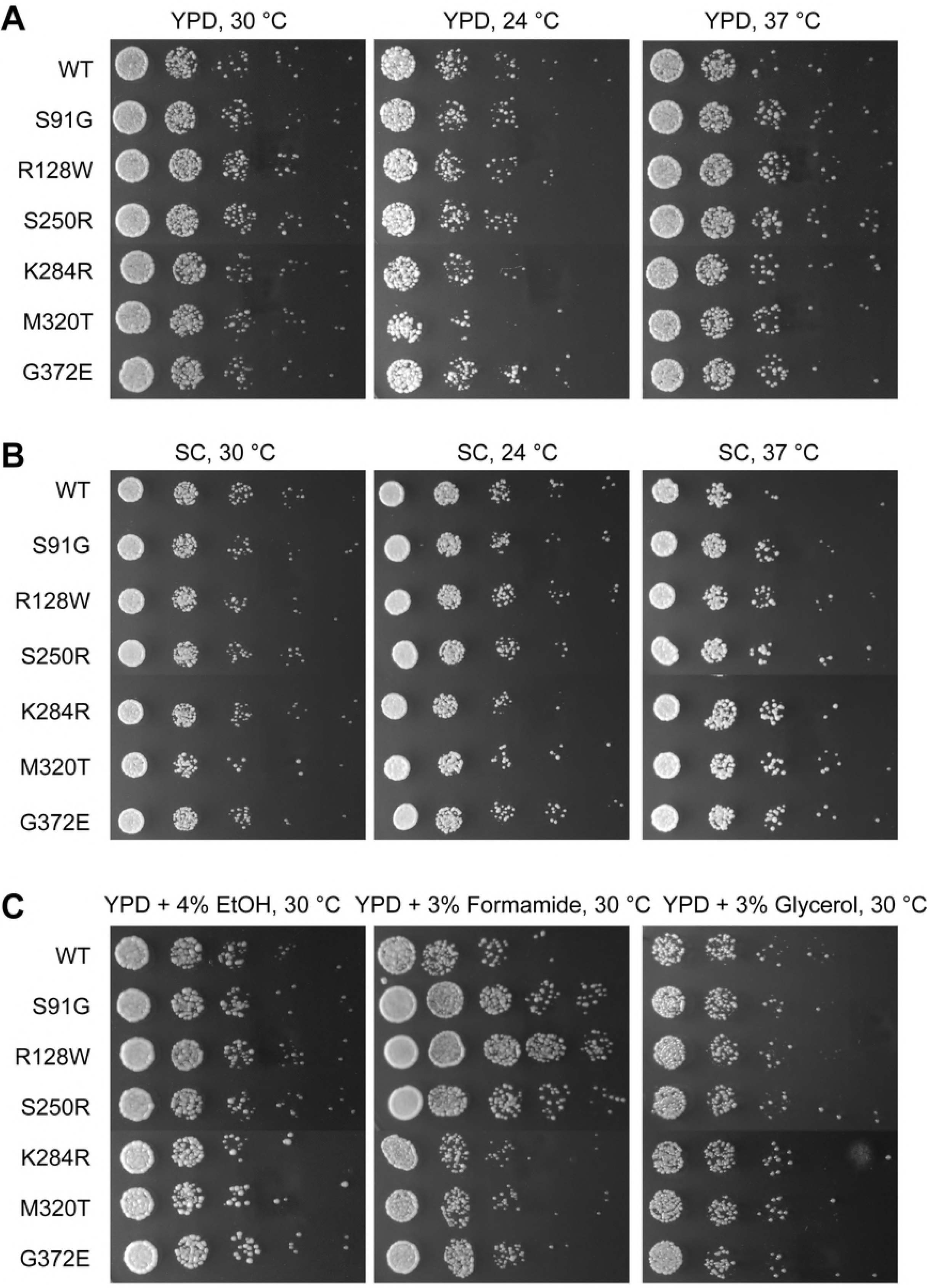
Growth curves of the wild-type and the mutant *cbf5* strains at different temperatures in liquid rich medium. Growth curves for BY4741 wild-type yeast are shown in comparison to mutant strains S91G, R128W and S250R (panels A, C and E) or strains K284R, M320T or G372E (panels B, D and F). Growth curves were recorded in a plate reader with shaking at the following temperatures: 30 °C (A and B), 18 °C (C and D) and 37 °C (E and F).

### Analyzing growth of *cbf5* mutant cells under stress conditions

Next, various stress conditions were tested for the presence of a phenotype of the mutant yeast cells using spot tests on solid agar media. First, temperature was varied to confirm the results observed in liquid medium (Figs 3A and B). Secondly, ethanol, formamide or glycerol was added to either rich medium (YPD) or synthetic complete (SC) medium (Fig 3C). Minimal medium was sometimes used in addition to rich medium, as it has been shown to enhance any milder phenotypes [25]. We also tested if mutations in *cbf5* result in a phenotype on plates containing 5-fluorouracil (Fig 4).

**Fig 3:**
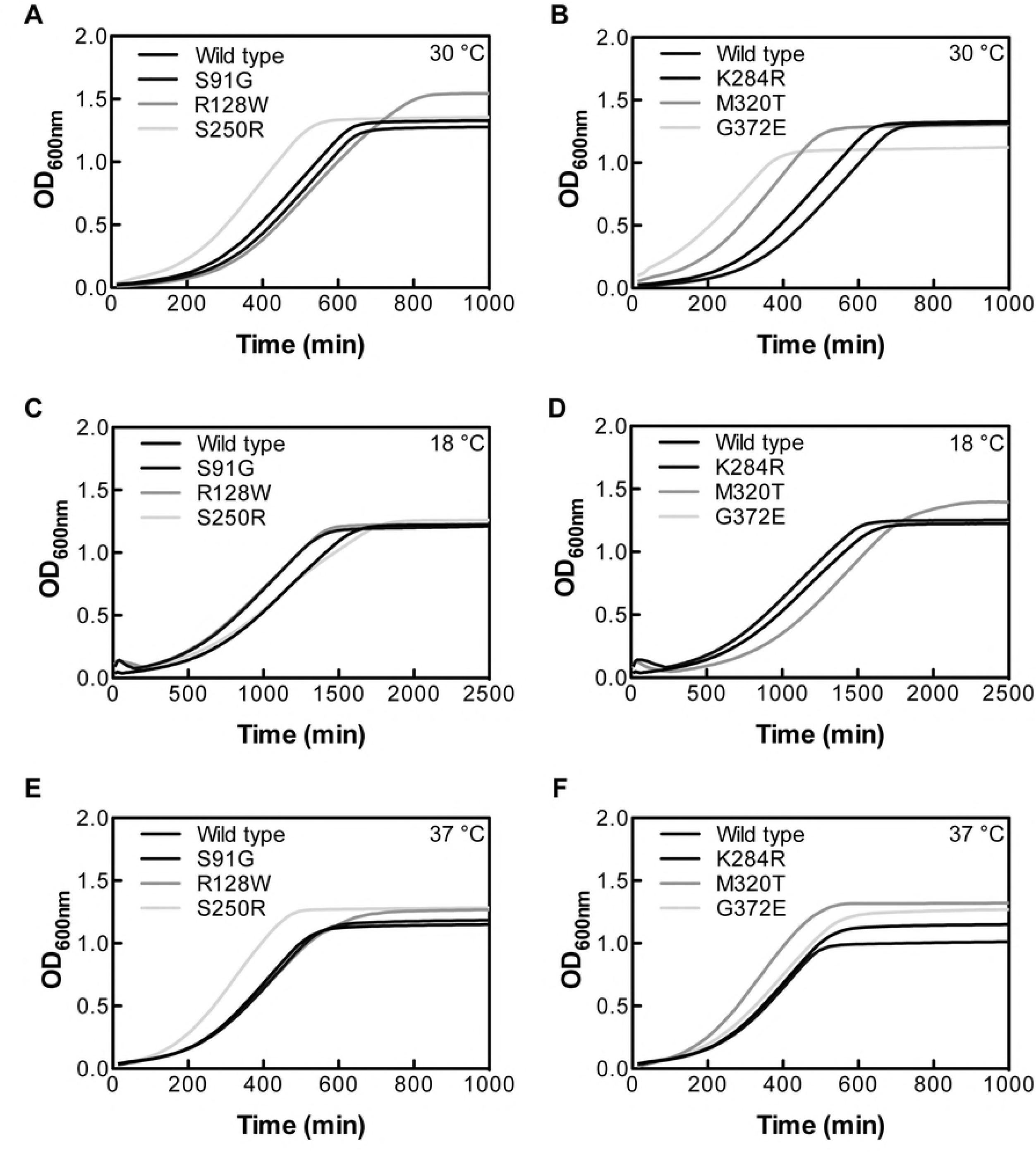
Spot tests of yeast strains with *cbf5* mutants under various stress conditions. Wild-type (BY4741) and mutant strains were grown at 24, 30 and 37 °C on rich media plates (A). The strains were also assessed for growth on minimal medium under the same temperature conditions (B). Lastly, cells were grown on YPD plates containing either 4% ethanol, 3% formamide or 3% glycerol at 30 °C (C).

**Fig 4:**
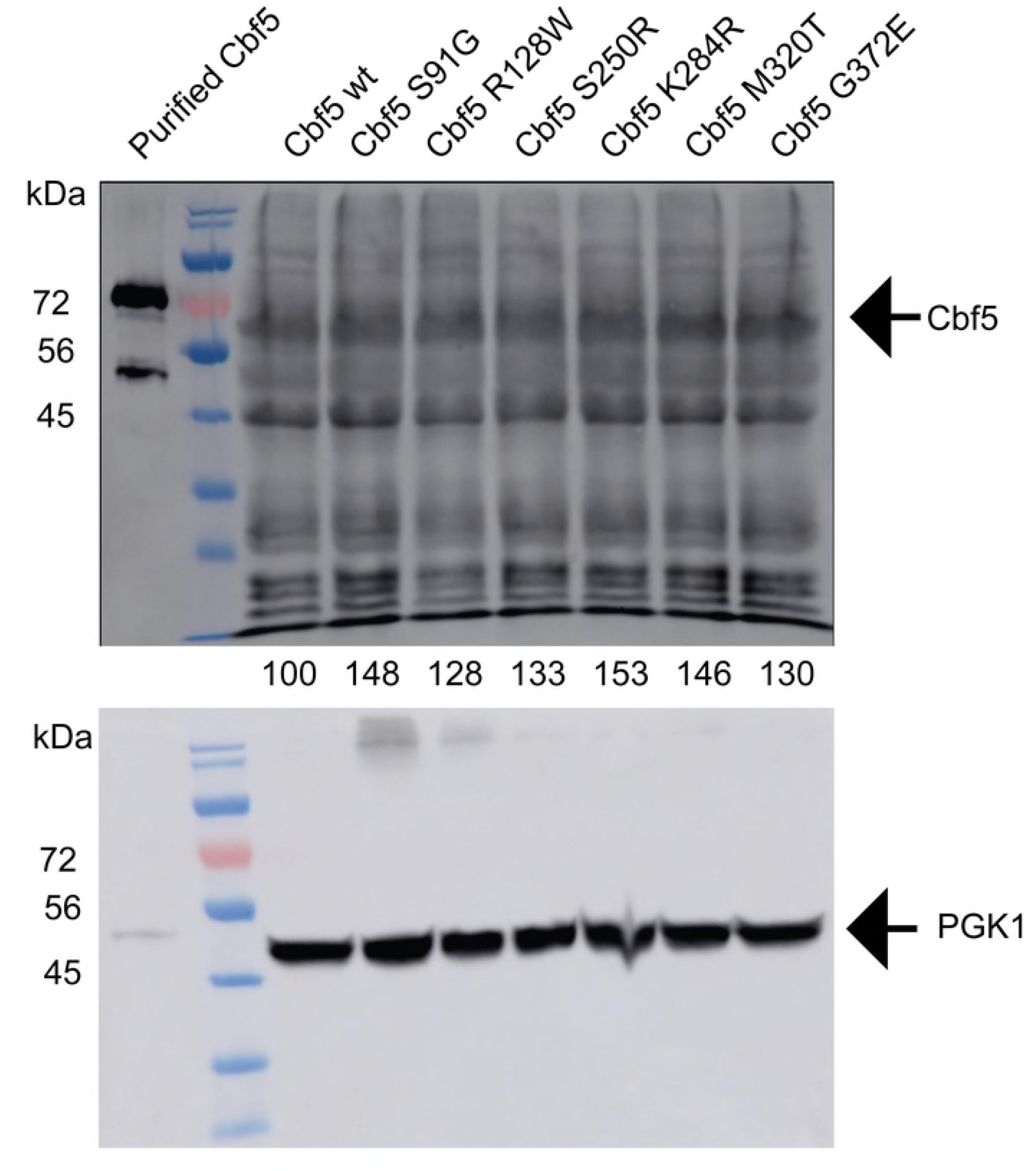
Spot tests in the presence of 5-fluorouracil. (A and B) An optimal 5-fluorouracil concentration was determined in both YPD (Panel A) or SC (Panel B) medium using BY4741 wild-type and the *cbf5* S91G strain. (C) Wild-type BY7471 and mutant strains were spotted onto SC plates containing 50 μM 5-fluorouracil which were grown at 30 °C.

Heat and cold sensitivity was tested by spotting mutant *cbf5* yeast cells onto both YPD and SC+ura plates followed by incubation at 24, 30 and 37 °C. As shown in Fig 3, all mutant strains grew equally well compared to wild-type at 30 °C on both YPD and SC+ura agar plates. Similarly, at a reduced temperature of 24 °C as well as an elevated temperature of 37 °C, no obvious phenotype was observed on either rich or minimal media for any of the *cbf5* mutant cells confirming the observations in liquid medium (Fig 2).

Next, ethanol was added to YPD at a concentration of 4% to examine the effects on protein stability since ethanol is a polar solvent and disrupts hydrogen bonds [25]. As shown in Fig 3C, ethanol had no effect on the growth of the mutant *cbf5* cells relative to wild-type. To further corroborate this observation, a 3% concentration of formamide was included in YPD medium to determine if the disruption of hydrogen bonds by a second polar solvent affected protein stability in the *cbf5* cellss. No growth defects were observed for any of the *cbf5* cells compared to wild-type cells suggesting the mutations to *cbf5* did not significantly reduce the stability of the protein (Fig 3C).

Furthermore, we assessed growth of the mutant cells on alternative carbon sources. When 3% glycerol was added to YPD medium, all mutant *cbf5* cells grew equally well when compared to the wild-type cells (Fig 3C). Since the mutant strains were able to use glycerol as a carbon source, this finding indicates that mitochondrial function remains unaffected when *cbf5* is mutated [25].

Pseudouridine synthases have been shown to be particularly sensitive to 5-fluorouracil, an inhibitor of pseudouridine formation [26–28]. Therefore, 5-fluorouracil was chosen to be included in the stress conditions that were tested. To first determine the concentration of 5-fluorouracil to use in yeast spot assays and whether YPD or SC medium would best show a phenotype, initial tests were performed in 6-well plates containing either 5 ml YPD or SC and a different 5-fluorouracil concentration ranging from 0 μM to 200 μM 5-fluorouracil [23]. Wild-type and *cbf5-S91G* cells were spotted in each well and allowed to grow at 30 °C. After 66 h of growth, 50 μM 5-fluorouracil inhibited the growth of both wild-type and *cbf5-S91G* cells to some extent while still forming colonies (Fig. 4 A and B). SC medium was chosen over YPD since it can enhance the appearance of a phenotype. Therefore, in subsequent yeast spot assays, a 5-fluorouracil concentration of 50 μM in SC medium was used. To ensure the DMSO used to dissolve the 5-FU was not affecting cell growth, 0.1% DMSO was added to control plates lacking 5-fluorouracil. All *cbf5* mutant cells grew as well as wild-type cells on plates containing 50 μM 5-fluorouracil (Fig 4C).

### Assessing Cbf5 protein levels

To ensure the substituted Cbf5 proteins were all being expressed to a similar level compared to wild-type Cbf5 protein, Western blotting was performed. Whole cell extract was prepared from each of the *cbf5* mutant as well as wild-type cells using equal amounts of cells. Cbf5 protein levels were determined using a Cbf5-specific antibody against 15 amino acids in the C-terminal region of Cbf5. In addition, primary antibody for phosphoglycerate kinase 1 (PGK1) was used as a loading control. Relative protein levels of each of the *cbf5* mutants cells indicate that the proteins were all being expressed to a similar level (Fig 5).

**Fig 5:**
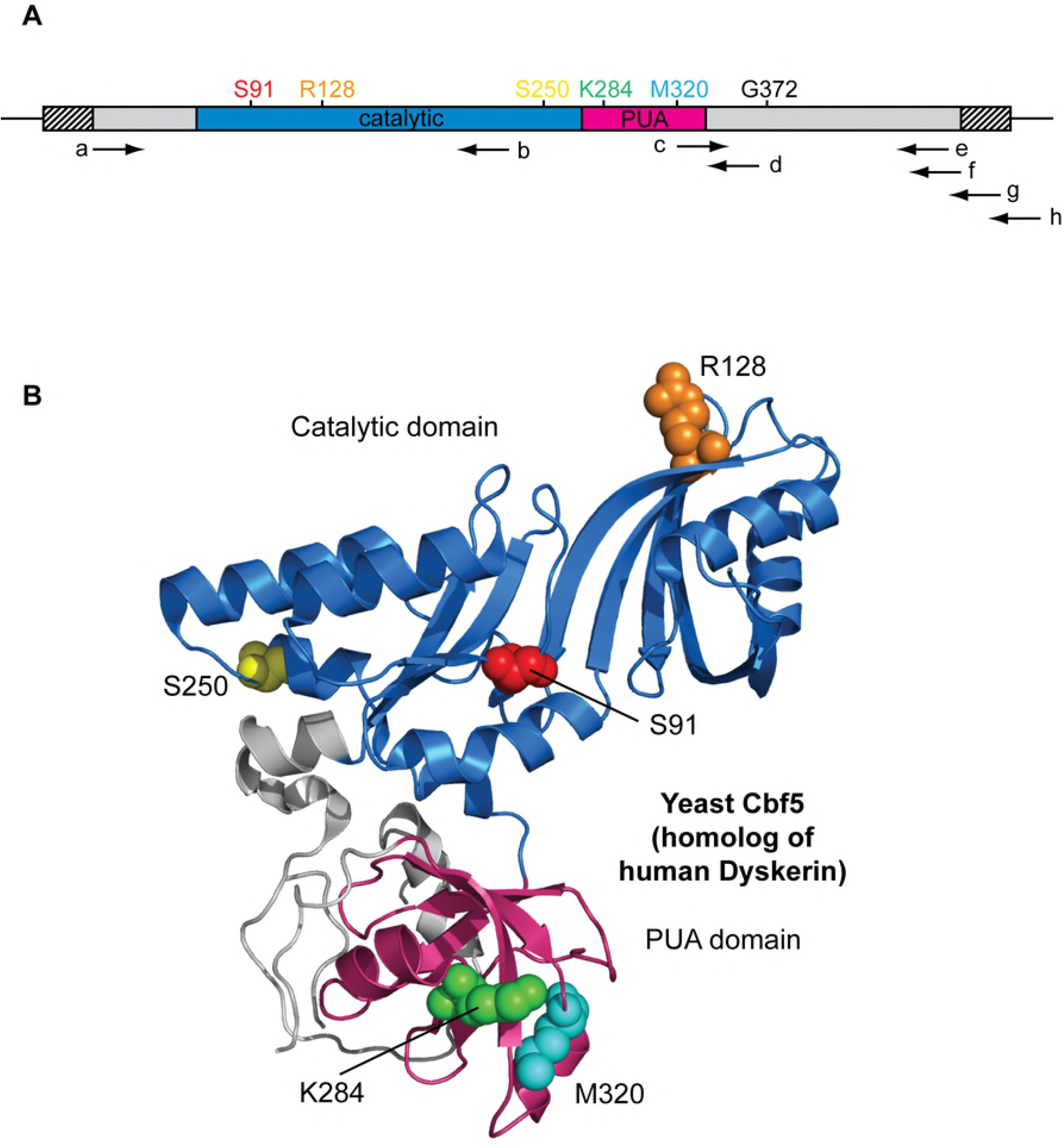
Western blot of Cbf5 from wild-type and mutant yeast strains. For Western blotting, a primary antibody against a small peptide chain in the C-terminus of Cbf5 was generated whereas PGK1 was detected as a loading control. S. cerevisiae Cbf5 is predicted to be 55 kDa in size, but presumably migrates a bit slower than expected due to post-translational modifications. Relative intensity of Cbf5 protein bands is indicated beneath each band. As a control (left lane), recombinantly expressed and purified ScCbf5 protein was analyzed [29]; this recombinant Cbf5 protein contains a hexahistidine tag and is likely differently modified than endogenous *S. cerevisiae* Cbf5 causing it to migrate differently.

## Discussion

To distinguish the effects of Dyskeratosis congenita mutations on telomerase stability and function compared to ribosome biogenesis, we created mutations in the *S. cerevisiae cbf5* gene corresponding to human disease mutations in the dyskerin gene because in yeast the dyskerin homolog, Cbf5, is only involved in ribosome biogenesis, but not in telomere maintenance. Therefore, we hypothesized that all observed effects would reflect the role of Cbf5 in ribosome biogenesis. All Cbf5 variants were expressed at similar protein levels indicating that they were equally stable as the Cbf5 wild-type protein. However, several different phenotypic assays in liquid and on solid medium, at different temperatures and different stress conditions did not show any obvious phenotype of the mutant *cbf5* cellss. Therefore, we conclude that *S. cerevisiae* can tolerate the Dyskeratosis congenita mutations in *cbf5*.

What can explain the absence of phenotypic effects of the *cbf5* mutations in yeast? Obviously, fungi will likely be less affected by such mutations in *cbf5* compared to mammalian cells because telomerase will remain functional in mutant yeast strains. Moreover, our results suggest that also other cellular functions such as ribosome biogenesis are not severely affected by *cbf5* mutations in *S. cerevisiae*. However, we cannot conclude from our results that Dyskeratosis congenita mutations in human dyskerin do not affect ribosome formation. Rather, differences between the mammalian and the yeast system might explain why no effect on ribosome formation and hence cellular phenotype were observed for the S. cerevisiae *cbf5* mutations.

H/ACA ribonucleoproteins play two important roles during ribosome biogenesis. Most H/ACA snoRNAs guide the site-specific pseudouridylation of rRNA catalyzed by the protein Cbf5 [13]. In contrast, the only essential yeast H/ACA snoRNA snR30 is mediating the processing of 18S precursor rRNA (pre-rRNA) [30]. The absence of a phenotype for the *cbf5* strains suggests that pre-rRNA processing by the essential snR30 guide RNA is not significantly affected implying that the Cbf5 protein variants harboring single-residue substitutions can still sufficiently interact with snR30. Interestingly, impairments of pre-rRNA processing have been reported for a zebrafish Dyskeratosis congenita model system; however in this case, expression levels of H/ACA proteins were reduced rather than introducing specific mutations [31, 32]. Presumably, yeast and possibly mammalian cells harbor mechanisms to prioritize the essential processing of pre-rRNA to take place even under sub-optimal conditions, e.g. when *cbf5* is mutated, but not absent.

It is noteworthy that we did not observe any reduction in protein levels for Cbf5 variants in the mutant yeast strains as vertebrate Dyskeratosis congenita models have been designed based on reduced Dyskerin expression [32–34]. However, the protein levels in Dyskeratosis congenita patients are not known. Most mutations in *dyskerin* lead to single-amino acid substitutions as assessed in this study whereas only a few mutations reside in introns and could potentially cause altered splicing and reduced protein levels [10]. We therefore speculate that most Dyskeratosis congenita mutations do not significantly destabilize the Dyskerin protein and decrease its level, but rather impair the function of Dyskerin.

The majority of H/ACA RNPs are responsible for rRNA modification by directing pseudouridine formation. Interestingly, the human ribosome harbors many more pseudouridines than the yeast ribosome, namely about 100 compared to 30 pseudouridines, respectively. Therefore, it is possible that the human ribosome depends much more on pseudouridine formation by H/ACA RNPs than the yeast ribosome. Indeed, yeast strains expressing only catalytically inactive Cbf5 protein that can form an RNP, but cannot form pseudouridines, display a strong cold- and heat-sensitive phenotype, but are viable suggesting that a yeast ribosome lacking pseudouridines can sustain cell growth [16]. Interestingly, studies in mammalian systems mimicking Dyskeratosis congenita have reported effects on translation, in particular internal ribosome entry site (IRES) mediated translation [33, 35, 36]. In yeast, ribosomes lacking pseudouridines show reduced fidelity and lower affinities for tRNAs and IRES elements [37]. Together, these studies suggest that Dyskeratosis congenita mutations affect translation only of a subset of mRNAs harboring IRES elements, but not necessarily all mRNAs. It is not clear how many mRNAs with IRES elements are used in fungi [38]. Hence, the impairment of IRES-mediated translation caused by *cbf5* mutations in yeast may affect only a small number of mRNAs and may therefore not result in an observable phenotype in contrast to human cells which might harbor many more IRES-dependent mRNAs.

Taken together, the lack of phenotypes observed for yeast strains harboring *cbf5* mutations suggests that yeast is not a good model system to study Dyskeratosis congenita. Both ribosome formation and ribosome function seem to be rather robust in *S. cerevisiae* and can tolerate single-residue substitutions in the Cbf5 protein. Similarly, deletions of most other pseudouridine synthases, that act in a guide-RNA-independent manner and modify tRNAs, snRNAs and mRNAs, do not cause strong phenotypes in yeast [39]. In general, the lack of phenotypes upon mutating *cbf5* or deleting other pseudouridine synthases might reflect the adaptation of single-cell fungi to unpredictable stress conditions requiring a robust translation apparatus in contrast to multi-cellular organisms where ribosome biogenesis and translation are exposed to less variable conditions and translation regulation is much more fine-tuned, e.g. through IRES-mediated mechanisms.

## Acknowledgements

We thank Dr. Linda Reha-Krantz for generous technical advice in creating the mutant yeast strains.

